# TargetSeeker-MS: A Computational Method for Drug Target Discovery using Protein Separation Coupled to Mass Spectrometry

**DOI:** 10.1101/513663

**Authors:** Mathieu Lavallée-Adam, Alexander Pelletier, Jolene K. Diedrich, William Low, Antonio F. M. Pinto, Salvador Martínez-Bartolomé, Michael Petrascheck, James J. Moresco, John R. Yates

## Abstract

When coupled to mass spectrometry (MS), energetics-based protein separation (EBPS) techniques, such as thermal shift assay, have shown great potential to identify the targets of a drug on a proteome scale. Nevertheless, the computational analyses assessing the confidence of drug target predictions made by these methods have remained rudimentary and significantly differ depending on the protocol used to produce the data. To identify drug targets in datasets produced using different EBPS-MS techniques, we have developed a novel flexible computational approach named TargetSeeker-MS. We showed that TargetSeeker-MS reproducibly identifies known and novel drug targets in *C. elegans* and HEK293 samples that were treated with the fungicide benomyl and processed using two different EBPS techniques. We also validated a novel benomyl target in vitro. TargetSeeker-MS, which is available online, allows for the confident identification of targets of a drug on a proteome scale, thereby facilitating the evaluation of its clinical viability.

## INTRODUCTION

Methods that can identify the putative protein targets of a drug on a proteome-scale are critical to decipher the mechanism of action of a compound. Such methods include large-scale phenotypic screenings, which are often performed to evaluate the ability of a library of compounds to modulate a given target pathway and therefore provide a putative treatment for the associated disease process. High-throughput screening (HTS) methods, which use automated high-end instrumentation, can test the properties of up to 100,000 compounds per day. However, high-throughput screening is typically performed on isolated systems and offers minimal insights regarding secondary protein targets^1^.

Mass spectrometry (MS)-based proteomics allows the large-scale identification and quantification of proteins in complex samples (e.g. cells, tissues, plasma). MS coupled to affinity-based enrichment strategies has routinely been used to identify protein interactions of compounds in the cell^2–4^. In the recent years, energetics-based protein separation (EBPS) techniques coupled to MS have emerged as a large-scale approach to rapidly and unbiasedly identify the protein targets of a given compound or drug. Such approaches rely on the hypothesis that the target of a drug will see its stability or thermodynamic properties changed upon binding with the compound. We previously demonstrated that Stability of Proteins from Rates of Oxidation (SPROX) combined with quantitative MS can be used to identify the protein targets of a drug^5^. This strategy uses the chemical denaturant-dependent oxidation rates of methionine residues to measure the thermodynamics of the unfolding or refolding reaction of proteins in drug-treated and untreated samples. A significant change in a given protein’s thermodynamics would indicate its binding to the drug. In this workflow, proteins were quantified using 6-plex tandem mass tags (TMT)^6^ stable isotope labeling coupled to MS. An increasingly popular EBPS coupled to MS (EBPS-MS) approach assesses ligand binding by evaluating changes in thermal stability^7,8^. This approach is based on the hypothesis that a protein bound by a given drug would see its thermal stability changed by the binding. A thermal shift assay technique was also employed in cells (CETSA)^9^. Savitski et al. demonstrated that combining a thermal shift assay approach with quantitative mass spectrometry allows the large-scale unbiased identification of drug targets^10^. These studies showed that EBPS-MS is complementary to HTS. Indeed, while HTS can process tens of thousands of compounds simultaneously and EBPS-MS only investigates one compound at a time, EBPS-MS allows the testing of this compound against the proteome of a sample, a much more complex system than what is tested in traditional HTS.

While EBPS-MS-based techniques for drug target discoveries are emerging rapidly, no general computational frameworks for the unbiased identification of drug-protein interactions have been produced. One of the reasons why the adoption of EBPS-MS technologies has been slow is because there are currently no implementations available to assess the confidence or statistical significance that a protein is bound by a given drug based on its change in stability evaluated by an EBPS-MS approach. The current practices involve the use of in-house computational scripts with numerous custom thresholds, making the comparisons and benchmarking of results across different laboratories extremely difficult^5,10^. Savitski et al. presented a statistical approach using curve fitting and statistics, which rely on largely unsupported assumptions of the normality distribution of the data^11^. In addition, this statistical approach is tied to the thermal shift assay experimental protocol using a 10-plex tandem mass tag as protein quantification technique^10^ and is not readily applicable to the other protein EBPS-MS quantification techniques that may be used to identify drug targets. For instance, we recently developed DiffPOP^12^, a novel EBPS technique that allows efficient separation of complex protein samples using an increasing concentration of a solution of acetic acid and methanol. DiffPOP was recently used to identify the targets of JIB-04, a compound that blocks the expression and transactivation of HIV-1 Tat^12^. DiffPOP differentiates itself from the Savitski et al. thermal shift assay approach by using MS to quantify proteins precipitated in each fraction instead of proteins remaining in the supernatant. The statistical method proposed by Savitski et al., which relies on melting curves based on supernatant analysis, is therefore not applicable for such a technique nor the SPROX quantitative MS approach. Furthermore, the use of TMT 10-plex reagents in Savitski et al.’s quantitative proteomics analysis suffers from a number of drawbacks. As in any TMT labeling analysis, low abundance proteins are less likely to be quantified, making it difficult to detect low abundance drug targets^13^. Furthermore, the price of TMT 10-plex reagents^14^ and the high resolution instruments necessary for TMT 10-plex analysis^15^ limits the democratization of Savitski et al.’s drug target discovery approach.

Herein, we propose a general computational framework, TargetSeeker-MS, for the identification of drug targets using EBPS coupled to quantitative MS. TargetSeeker-MS implements a Bayesian inference machine learning approach to assess the confidence that a protein is bound by a given compound. We demonstrate that TargetSeeker-MS, which is open-source and available as a user-friendly web-server, is hypothesis-free and flexible enough to analyze datasets originating from any EBPS-MS techniques. TargetSeeker-MS identified putative targets of benomyl, a fungicide putatively linked to Parkinson’s disease^16,17^, in two *C. elegans* datasets analyzed using DiffPOP and thermal shift assay coupled to MS. We showed that although both fractionation methods vary in nature, TargetSeeker-MS predictions in both datasets share a significant overlap of confident targets. In addition, we demonstrated that the TargetSeeker-MS algorithm predicts drug targets with a greater sensitivity than previously proposed approaches. Benomyl is known to inhibit aldehyde dehydrogenase (ALDH), a mechanism putatively leading to Parkinson’s disease development^16^. TargetSeeker-MS identified aldehyde dehydrogenase as a benomyl target along with other known and novel targets. Finally, we highlight that TargetSeeker-MS identified human benomyl target orthologs when processing a HEK293 cells dataset treated with the drug and validated the impact of benomyl on the enzymatic activity of one of its novel predicted targets, GAPDH.

## RESULTS

TargetSeeker-MS is a Bayesian inference-based approach that computes the probability that a protein is bound by a given drug through the analysis of EBPS-MS datasets. Briefly, TargetSeeker-MS takes as input a set of untreated (control) samples that were processed using EBPS and quantified using MS and builds for each protein a noise model of the similarity of the protein fractionation profiles in different biological replicates. It then evaluates the similarity of these control protein fractionation profiles with that of a drug-treated sample that was also separated using the same EBPS-MS approach. TargetSeeker-MS then assesses the confidence that each protein is bound by the drug. Figure 1 provides a graphical representation of TargetSeeker-MS’ pipeline. In this study, we used TargetSeeker-MS to identify the proteins bound by benomyl in three different datasets. The first two datasets analyzed *C. elegans* samples and were produced using DiffPOP separation (see Methods) coupled to MS (Dataset 1 – DiffPOP/*C. elegans*; Supplementary Figure S1) and Thermal Shift Assay (TSA) separation coupled to MS (Dataset 2 – TSA/*C. elegans*; Supplementary Figure S1). The third dataset represents the processing of Human Embryonic Kidney 293 cells (HEK 293) using DiffPOP-MS (Dataset 3 – DiffPOP/HEK293; Supplementary Figure S1).

**Figure 1.**
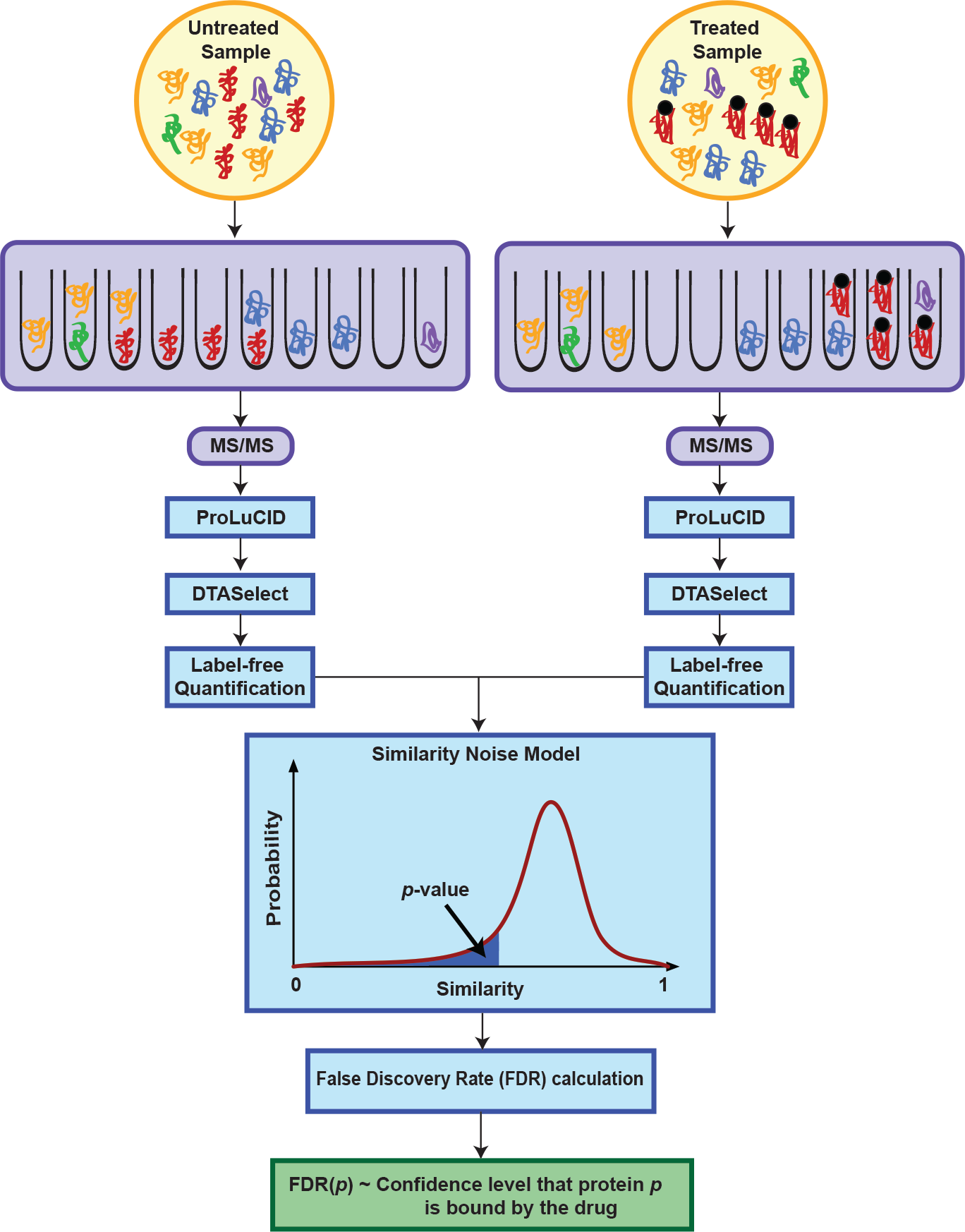
Experimental and computational pipeline. A graphical representation of the drug target identification pipeline illustrating the protein separation, the mass spectrometry analysis, and the TargetSeeker-MS algorithm.

### Protein fractionation profiles display a high level of reproducibility

To begin, we investigated the level of reproducibility of protein (*p*) fractionation profiles 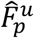 across different biological replicates of untreated biological samples (*u*) in order to assess the possibility of building a model of the noise of fractionation profile similarity values with a small number of replicates. With this objective in mind, we computed the similarity 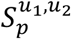 for all proteins *p* and all untreated samples *u*_1_ and *u*_2_ ∈ *U*, where *U* is the set of untreated samples in a dataset. The vast majority of the protein fractionation profiles shared a high degree of similarity (Figure 2A and Supplementary Figures S2 and S3). For instance, 79% of all pairs of fractionation profiles in untreated samples have a similarity 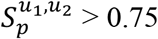 in the DiffPOP-*C. elegans* dataset (Dataset 1). These similarity values demonstrate that both EBPS-MS approaches yield a high level of fractionation reproducibility, which is likely to be sufficient to compute the probability matrix 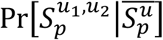 and to build an accurate noise model of the similarity values of fractionation profiles under the null hypothesis (untreated samples) with a small number of replicates (see Methods). Supplementary Figure S4 displays the probability matrix 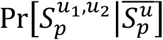 for Dataset 1. Protein fractionation profiles also shared a high similarity in benomyl treated samples (Supplementary Figure S5, S6, and S7). To illustrate this, in Dataset 1 83% of all pairs of fractionation profiles obtained a similarity 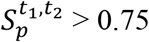, where *t*_1_ and *t*_2_ ∈ *T* the set of all treated samples. The similarity values between the untreated and benomyl treated samples are similarly distributed in all three datasets with only the DiffPOP-HEK293 dataset (Dataset 3) showing a slight difference between the two distributions (Figure 2B, Supplementary Figure S8, and S9). Overall, the distributions of similarity values highlight the feasibility of creating a noise model for the similarity between the fractionation profiles of a given protein using a small number of biological replicates.

**Figure 2.**
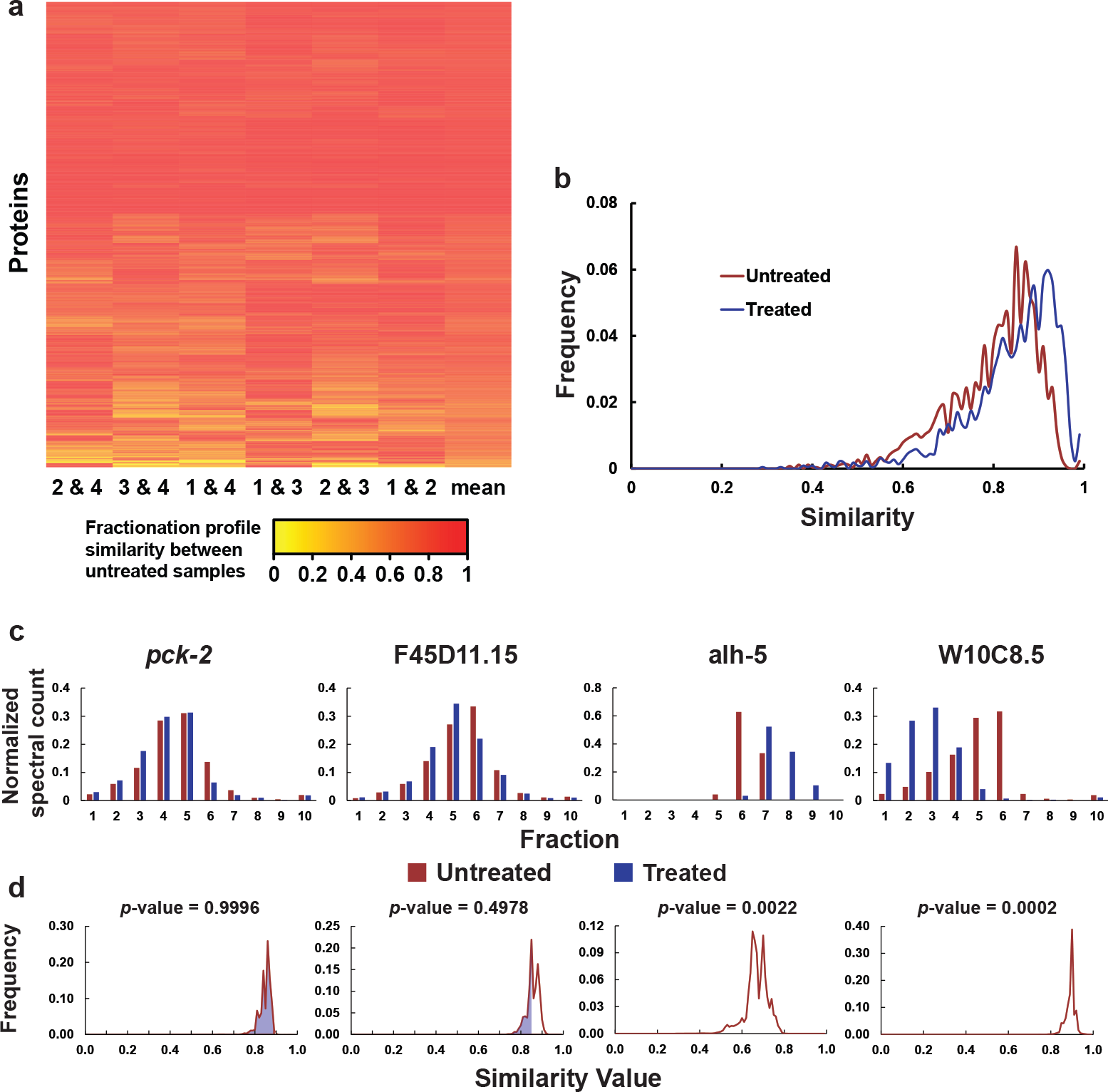
Fractionation profile similarity. (A) Heatmap representation of the similarity between protein fractionation profiles for all pairs of untreated samples of the DiffPOP/ *C. elegans* dataset. All proteins with a sufficient abundance to compute a fractionation profile in all untreated samples are displayed. (B) Distributions of the fractionation profile similarity values in both untreated and benomyl treated samples of the DiffPOP/*C. elegans* dataset. (C) Fractionation profiles represented as normalized spectral counts in each fraction and (D) posterior distributions of 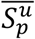 of four different C. *elegans* proteins identified using DiffPOP-MS. The shaded portion of the distributions represents the *p*-value 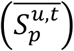 for each protein. The shaded portion of W10C8.5 and alh-5 is not visible since it spans from (0 to 0.62) and (0 to 0.38), respectively.

### TargetSeeker-MS assesses the statistical significance of protein fractionation profile changes upon benomyl treatment

The ability of the algorithm to assess the significance of the change in the fractionation profile of a protein upon drug treatment was illustrated with four protein examples from Dataset 1 (Figure 2C and 2D). *Pck-2* represents an example of a protein without a change in its fractionation profile upon benomyl treatment (Figure 2C). Indeed, both distributions of normalized spectral counts are almost identical (Similarity 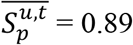). The associated *p*-value (>0.99; FDR=1.0) computed by TargetSeeker-MS is therefore very high (see Methods for *p*-value calculation). On the other hand, F45D11.15 appears to display a small shift to the left in its fractionation profile upon benomyl treatment (Similarity 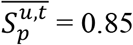). However, this change in fractionation profile is too minor to be deemed significant by TargetSeeker-MS (*p*-value = 0.5; FDR = 1.0), since it may simply be due to noise as indicated by its high FDR. *Alh-5*, a protein known to be bound by benomyl^16,18^ clearly displays a significant change in its fractionation profile, showing an increased precipitation resistance upon benomyl treatment (Similarity 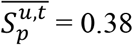). Due to the amplitude of this shift, TargetSeeker-MS assigned a *p*-value of 0.0022 (FDR=0.005) to *alh-5*. Conversely, W10C8.5, an ortholog of a human creatine kinase, also sees its fractionation profile drastically modified by benomyl (Similarity 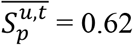), but in this case a decrease in precipitation resistance is observed (*p*-value = 0.0002; FDR< 0.005). This result is consistent with previous reports, which showed that creatine kinase enzymatic activity is altered by benomyl^19,20^.

### TargetSeeker-MS identifies high-confidence benomyl targets in *C. elegans* samples that were analyzed using DiffPOP-MS

We tested the ability of TargetSeeker-MS to identify benomyl targets in *C. elegans* through the analysis of a dataset produced with DiffPOP-MS (Dataset 1). TargetSeeker-MS built the similarity noise model with a set of four untreated biological replicates. With this noise model established, we used TargetSeeker-MS to evaluate the confidence that the precipitation resistance of proteins quantified in three biological replicates of benomyl treated *C. elegans* samples was altered. We first analyzed all drug treated samples as a group in a single TargetSeeker-MS analysis, computing the average of the similarity values between 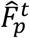 and 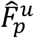 for all treated samples *t* and all untreated samples *u*. TargetSeeker-MS identified 59 proteins with a FDR < 0.01 and 101 with a FDR < 0.05 (Supplementary Table S1).

We benchmarked the TargetSeeker-MS analysis against two alternate approaches: the *Z*-score method and the Savitski et al. statistical approach (see Methods) (Figure 3A). Using high-confidence FDRs, TargetSeeker-MS reported more drug target predictions than either of the other two methods. To maximize the stringency of TargetSeeker-MS drug target predictions, in addition to the FDR threshold, a Fold-change of Similarity Difference (FSD) threshold of 0.20 was applied to each protein (see Methods). Using both thresholds, TargetSeeker-MS identified 41 proteins with a FDR < 0.05 and a FSD > 0.2 (Figure 3B).

**Figure 3.**
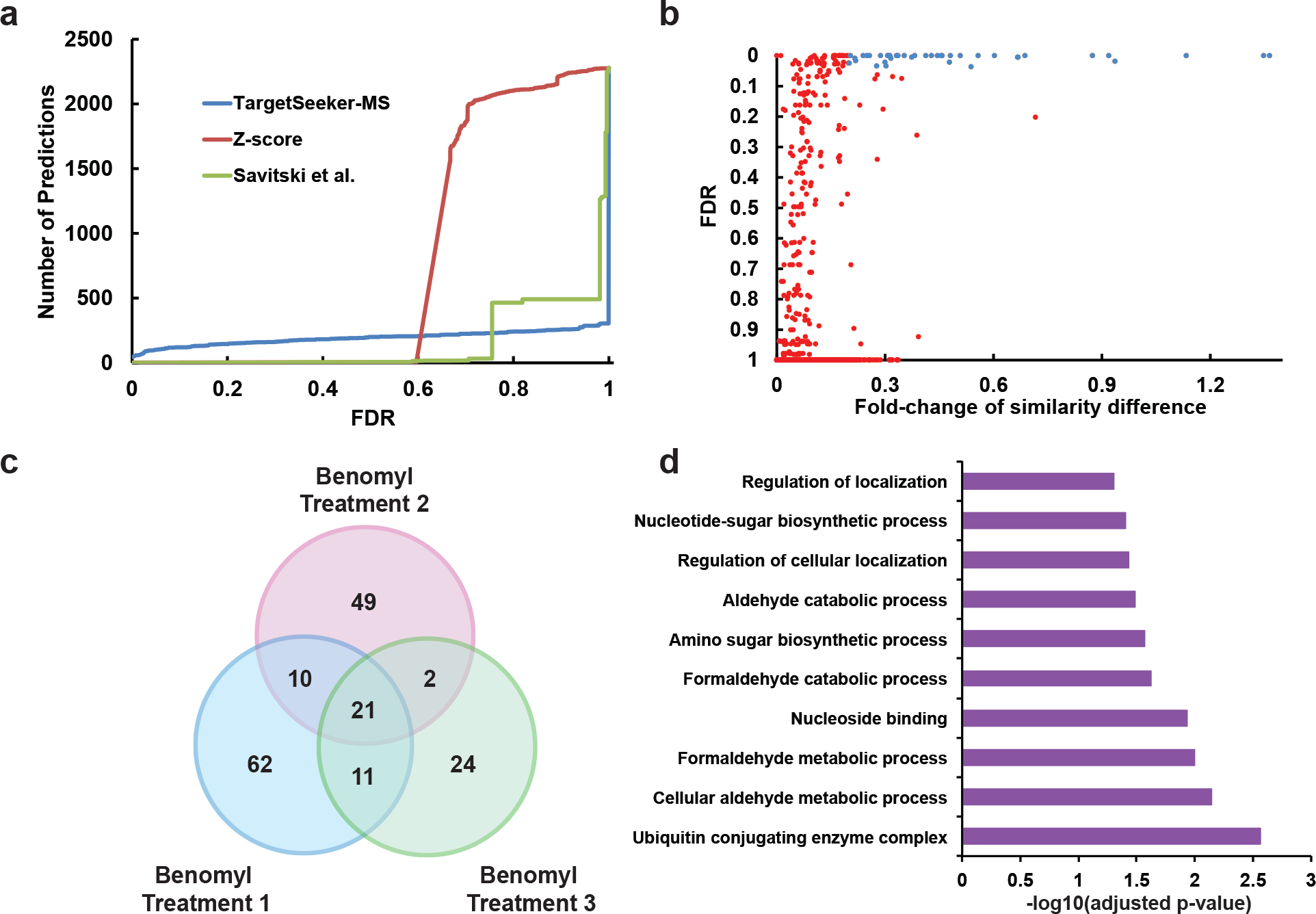
Analysis of the DiffPOP/*C. elegans* dataset. (A) Cumulative distributions of the number of target predictions at a given FDR obtained by TargetSeeker-MS, the Z-score method, and the Savitski et al. approach. (B) FDR and fold-change of similarity difference of all proteins to which a fractionation profile was assigned. High-confidence drug targets (FDR < 0.1, FSD > 0.2) are represented in blue, while low-confidence predictions are shown in red. (C) Venn diagram representation of the high-confidence predictions of TargetSeeker-MS when processing each drug treated biological replicate independently. (D) Gene Ontology terms that are enriched among the drug targets identified by TargetSeeker-MS in all three biological replicates.

While TargetSeeker-MS can assess the significance of the average of the similarity values between 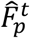 and 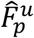 for all treated samples *t* and all untreated samples *u*, it can also assess the significance of the similarity between 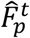 and 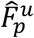 for all untreated samples and a single given treated sample *t*. This allows TargetSeeker-MS to independently identify drug targets in each treated sample. Proteins that are reproducibly predicted as drug targets in all treated samples (biological replicates) therefore represent very high confidence predictions. To once again maximize the stringency of our analysis, we only considered proteins as drug targets if they were identified as high-confidence targets by TargetSeeker-MS (FDR <0.1, FSD > 0.2) in all treated samples (Figure 3C; Supplementary Table S2-S5). The resulting 21 proteins are reported in Table 1. This list includes alh-5, an aldehyde dehydrogenase. It was previously reported that benomyl inhibits the low-Km hepatic mitochondrial aldehyde dehydrogenase of mice^18^ and that the inhibition of aldehyde dehydrogenase may lead to preferential development of Parkinson’s disease^16^. TargetSeeker-MS also identified a number of creatine kinases (F32B5.1, F44G3.2, W10C8.5, ZC434.8) as high-confidence benomyl targets. These results are consistent with studies demonstrating that the enzymatic activity of a creatine kinase was altered by benomyl in rats^19^ and in human serum^20^. Unc-25, a *C. elegans* ortholog of the GABA neurotransmitter biosynthetic enzyme that was not previously reported as a benomyl target, was also identified by TargetSeeker-MS to be affected by the compound. The 21 targets identified by TargetSeeker-MS are significantly enriched for Gene Ontology (GO) terms (functions and biological processes) that have been reported to be affected by benomyl, such as “Aldehyde catabolic process” and “Cellular aldehyde metabolic process” (Figure 3D; Supplementary Table S6 for complete enrichment analysis results)^16,18^.

**Table 1:** Benomyl targets as predicted by TargetSeeker-MS in the *C. elegans*/DiffPOP dataset.^1^.

| Wormbase ID | Gene Name<sup>2</sup> | Untreated Similarity | Drug vs Untreated Similarity | p-value | FDR | Fold-change of similarity difference |
| --- | --- | --- | --- | --- | --- | --- |
| F44G3.2 | CKB*, CKM*, CKMT2*, CKMT1B* | 0.90 | 0.70 | $3.2 \times 10^{-5}$ | <0.005 | 0.29 |
| Y54G2A.31 | ubc-13 | 0.90 | 0.65 | $3.4 \times 10^{-5}$ | <0.005 | 0.38 |
| F20G2.2 | RDH8, DHRS1 * | 0.85 | 0.56 | $8.0 \times 10^{-5}$ | <0.005 | 0.51 |
| T07C4.9b | nex-2 | 0.87 | 0.60 | $2.0 \times 10^{-4}$ | <0.005 | 0.44 |
| T22D1.3a | IMPDH2*, IMPDH1* | 0.81 | 0.55 | $2.0 \times 10^{-4}$ | <0.005 | 0.48 |
| W10C8.5 | CKB*, CKMT2*, CKMT1B* | 0.90 | 0.62 | $2.0 \times 10^{-4}$ | <0.005 | 0.46 |
| Y37D8A.23a | unc-25 | 0.83 | 0.52 | $2.0 \times 10^{-4}$ | <0.005 | 0.60 |
| Y37D8A.23b | unc-25 | 0.82 | 0.52 | $2.0 \times 10^{-4}$ | <0.005 | 0.56 |
| ZC434.8 | CKB*, CKMT2*, CKMT1B* | 0.89 | 0.71 | $2.0 \times 10^{-4}$ | <0.005 | 0.25 |
| F32B5.1 | CKB*, CKM*, CKMT2*, CKMT1B* | 0.89 | 0.61 | $4.0 \times 10^{-4}$ | <0.005 | 0.46 |
| T22D1.3b | IMPDH2*, IMPDH1* | 0.77 | 0.45 | $5.0 \times 10^{-4}$ | <0.005 | 0.69 |
| Y42G9A.4c | mvk-1 | 0.82 | 0.63 | $1.1 \times 10^{-3}$ | <0.005 | 0.31 |
| C36A4.4 | UAP1*, UAP1L1* | 0.81 | 0.57 | $1.2 \times 10^{-3}$ | <0.005 | 0.43 |
| Y37D8A.23c | unc-25 | 0.67 | 0.36 | $1.2 \times 10^{-3}$ | <0.005 | 0.87 |
| K06B9.2 | UAP1*, UAP1L1* | 0.80 | 0.57 | $1.5 \times 10^{-3}$ | <0.005 | 0.41 |
| Y48G10A.1 | ESD* | 0.73 | 0.44 | $1.9 \times 10^{-3}$ | 0.005 | 0.67 |
| T08B1.3 | alh-5 | 0.64 | 0.38 | $2.2 \times 10^{-3}$ | 0.005 | 0.67 |
| F21C3.3 | hint-1 | 0.54 | 0.28 | $5.4 \times 10^{-3}$ | 0.019 | 0.94 |
| D2063.3a | glrx-3 | 0.78 | 0.60 | $6.0 \times 10^{-3}$ | 0.021 | 0.30 |
| R07E3.1a | CTSF | 0.72 | 0.49 | $6.2 \times 10^{-3}$ | 0.021 | 0.48 |
| D2063.3b | glrx-3 | 0.77 | 0.59 | $9.9 \times 10^{-3}$ | 0.034 | 0.30 |

<sup>1</sup> A yellow row represents a protein that was not quantified with enough confidence in the TSA/*C. elegans* dataset to allow TargetSeeker-MS to assess the significance of benomyl binding. A green row represents a protein that was also predicted as a benomyl target in the TSA/*C. elegans*.
<sup>2</sup> A \* indicates the human ortholog gene name of a *C. elegans* protein without a name.

### TargetSeeker-MS identified high confidence benomyl targets in the TSA/*C. elegans* dataset

In order to test the ability of the TargetSeeker-MS algorithm to accurately identify drug targets in samples that were separated using a different EBPS-MS method, we applied our algorithm to the TSA-*C. elegans* dataset (Dataset 2). Combining two replicate TSA analyses of benomyl-treated samples into a single TargetSeeker-MS analysis allowed the algorithm to identify 278 benomyl targets with a FDR < 0.01 and 331 benomyl targets with a FDR < 0.05 (Supplementary Table S7).

Even though Dataset 2 was processed using a similar approach to that described in Savitski et al. (i.e. using the same mechanism: heat destabilization), TargetSeeker-MS again identified more drug targets at high-confidence FDRs than the Z-score approach or the Savitski et al. method (Figure 4A). Interestingly, 285 proteins in Dataset 2 that were assigned a FDR < 0.05, were associated with a relatively low FSD (< 0.2) (Figure 4B). This result is likely explained by the smaller variance in similarity values of fractionation profiles in replicate samples obtained with TSA-MS compared to DiffPOP-MS (Figure 2B and Supplementary Figure S8). This decreased variation in fractionation profiles therefore allows TargetSeeker-MS to assign high-confidence FDRs to even small changes in protein fractionation profiles upon drug treatment. To maximize sensitivity we therefore opted to not apply a FSD threshold for all data produced using TSA-MS. Nevertheless, in order to maintain a high level of confidence in TargetSeeker-MS predictions, we only retained as putative drug targets proteins that were assigned by the algorithm a FDR < 0.05 in both replicate TSA analyses of benomyl-treated samples when analyzed independently (Supplementary Table S8, S9, and S10). This process yielded a list of 154 benomyl-binding proteins in the TSA/*C. elegans* dataset. These proteins are statistically significantly enriched for GO terms such as “Microtubule” (adjusted *p*-value = 0.0075), “Regulation of cellular protein localization” (adjusted *p*-value = 0.043), “Structural constituent of cytoskeleton” (adjusted *p*-value = 0.0013), and “Cell periphery” (adjusted *p*-value = 0.033) (Figure 4C and Supplementary Table S11). These GO term enrichments are consistent with the role of benomyl, which was previously reported as a compound that depolymerizes microtubules near the cell periphery^21–23^.

**Figure 4.**
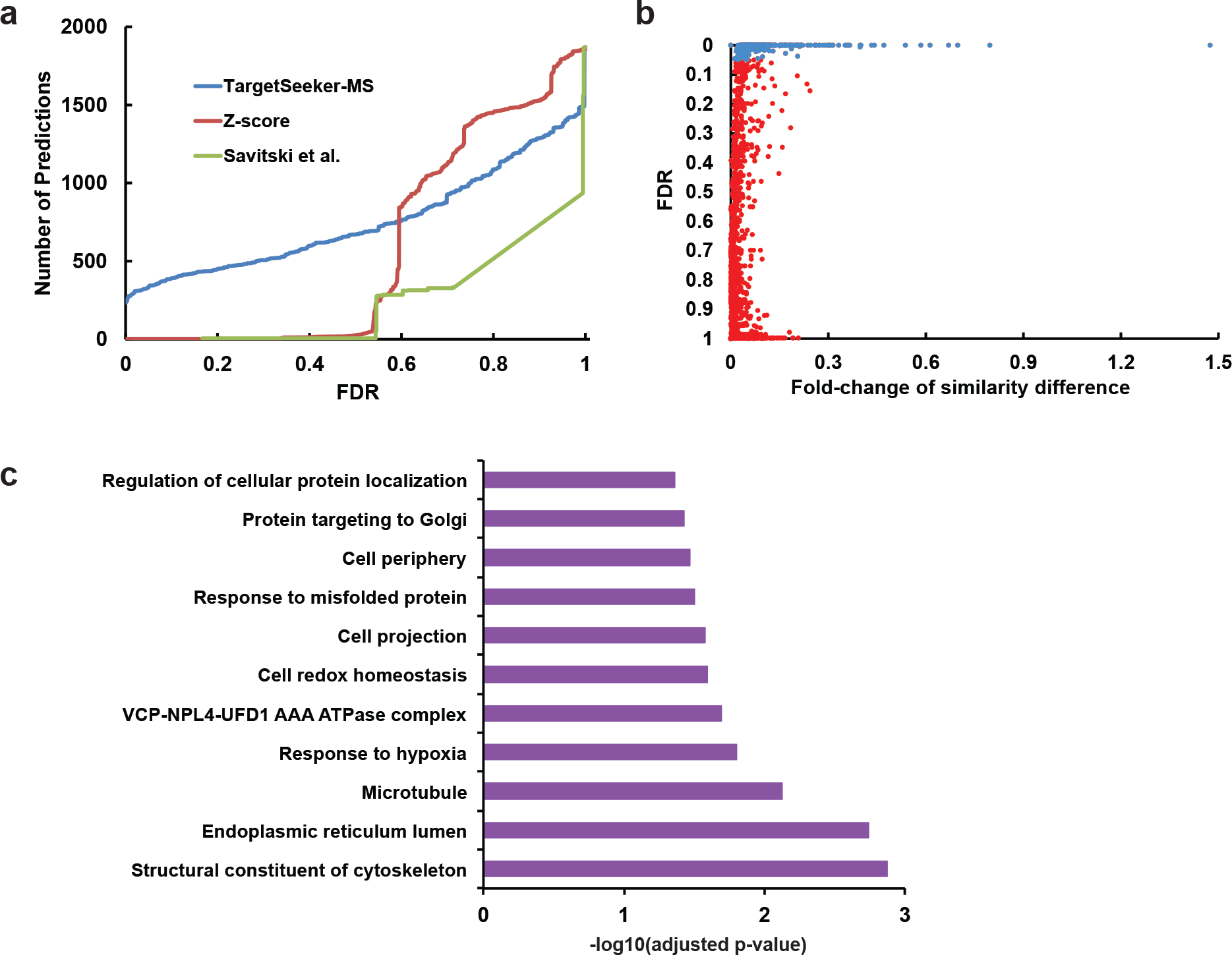
Analysis of the TSA/*C. elegans* dataset. (A) Cumulative distributions of the number of target predictions at a given FDR obtained by TargetSeeker-MS, the Z-score method, and the Savitski et al. approach. (B) FDR and fold-change of similarity difference of all proteins to which a fractionation profile was assigned. High-confidence drug targets (FDR < 0.05) are represented in blue, while low-confidence predictions shown in red. (C) Gene Ontology terms that are enriched among the drug targets identified by TargetSeeker-MS in both biological replicates.

### Drug targets identified in the TSA/*C. elegans* dataset significantly overlap with those identified in the DiffPOP/*C. elegans* dataset

Among the *C. elegans* benomyl targets identified in the TSA dataset, we count a number of creatine kinases (F44G3.2, W10C8.5, and F32B5.1) that were also identified in the DiffPOP-MS/*C. elegans* dataset. However, alh-5 a known target of benomyl was not quantified with enough spectral counts to be associated with a fractionation profile and could therefore not be assessed by TargetSeeker-MS (see Methods). Even though TSA and DiffPOP use different mechanisms to separate proteins, the intersection of the sets of drug targets identified by TargetSeeker-MS in both datasets is significant (hypergeometric test *p*-value = 0.02). Indeed, 8 out of the 11 drug targets from the DiffPOP/*C. elegans* that were quantified accurately enough in the TSA/*C. elegans* dataset to be analyzed by TargetSeeker-MS were also identified as drug targets in the latter dataset (Table 1). These results highlight the ability of TargetSeeker-MS to reproducibly detect protein targets in datasets originating from different EBPS-MS techniques.

### TargetSeeker-MS identifies high-confidence benomyl targets in HEK 293 that are *C. elegans* orthologs

We executed the TargetSeeker-MS algorithm on the DiffPOP/HEK293 datasets (Dataset 3) to obtain some insights regarding human proteins that may be bound by benomyl. TargetSeeker-MS identified 349 human drug targets with a FDR < 0.01 and 566 proteins with an FDR < 0.05 (Supplementary Table S12). Once again, TargetSeeker-MS outperformed the *Z*-score method and the Savitski et al. statistical approach by detecting more benomyl targets at high confidence FDRs (Figure 5A). To maximize the specificity of the analysis, a FSD threshold of 0.2 was applied yielding 77 high-confidence drug predictions with a FDR < 0.01 and 94 with a FDR < 0.05 (Figure 5B; Supplementary Table S12). To maximize the specificity of the drug predictions as for the other datasets, we only retain as putative drug targets proteins that were assigned by the algorithm a FDR < 0.05 in both benomyl-treated replicates when analyzed independently (Supplementary Table S13, S14, and S15). This resulted in a list of 22 high-confidence benomyl targets (Supplementary Table S15). Among these targets we find three proteins, CKMT1A (FDR < 0.001), CKMT2 (FDR < 0.001), and UBE2O (FDR = 0.005) that correspond to orthologs of *C. elegans* proteins (E-value < 10^−6^ and sequence identity > 50%; see Methods) that were also predicted by TargetSeeker-MS to be high-confidence benomyl targets in the DiffPOP/*C. elegans* dataset (Dataset 1). In addition, CKMT1A and CKMT2 orthologs were also identified as benomyl targets in the TSA/*C. elegans* dataset (Dataset 2). Of note, creatine kinases were identified as benomyl targets in all three datasets that involve two different EBPS-MS techniques analyzing two different organisms. Among the notable GO term enrichments present in the list of 22 benomyl targets, we find “Creatine metabolic process” and “Creatine kinase activity”, which are biological processes that have been reported to be linked to benomyl^19,20^ (Figure 5C and Supplementary Table S16).

**Figure 5.**
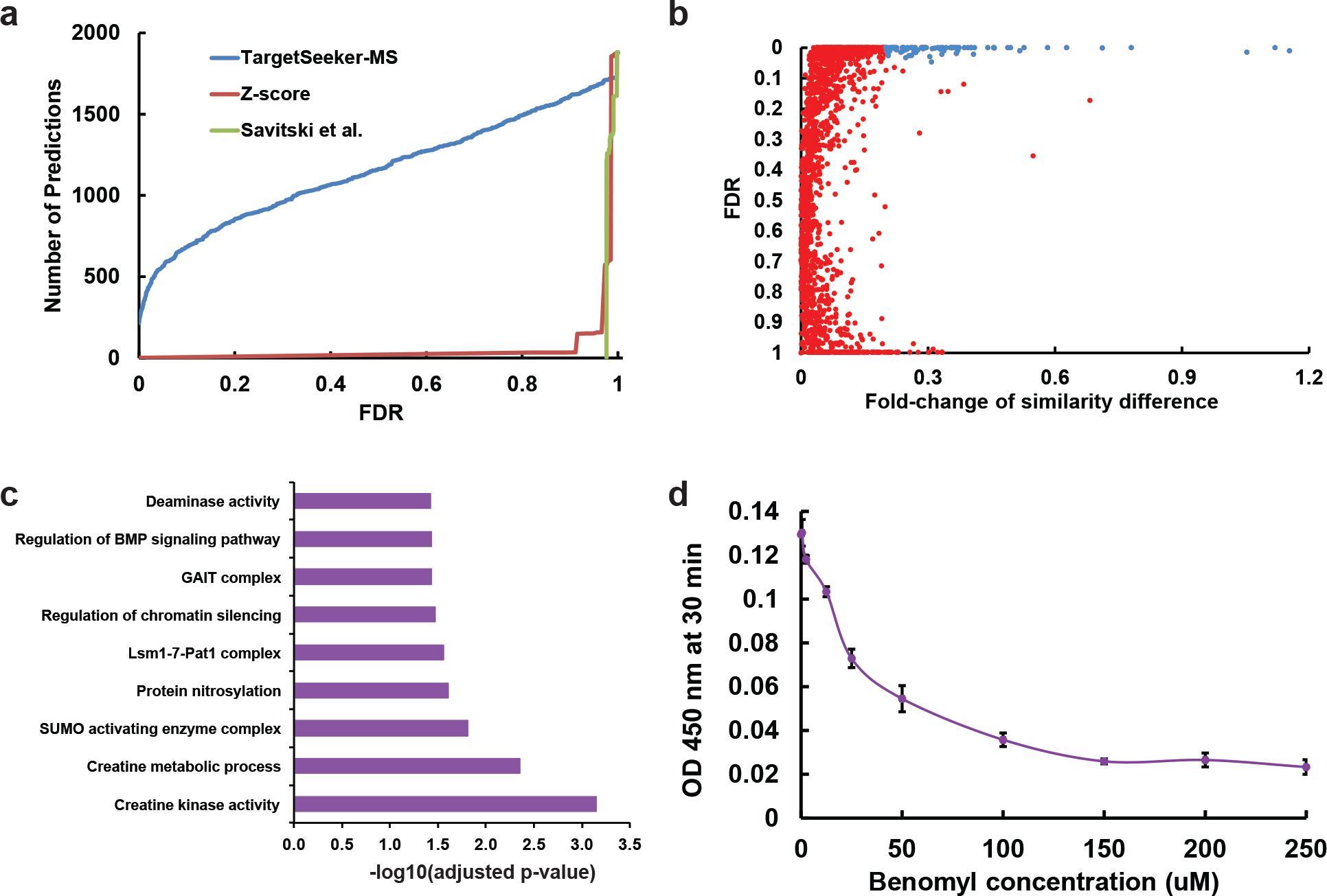
Analysis of the DiffPOP/HEK 293 dataset. (A) Cumulative distributions of the number of target predictions at a given FDR obtained by TargetSeeker-MS, the Z-score method, and the Savitski et al. approach. (B) FDR and fold-change of similarity difference of all proteins to which a fractionation profile was assigned. High-confidence drug targets (FDR < 0.05, FSD > 0.2) are represented in blue, while low-confidence predictions shown in red. (C) Gene Ontology terms that are enriched among the drug targets identified by TargetSeeker-MS in both biological replicates. (D) Efficiency of benomyl inhibition of GAPDH enzymatic activity upon mixing with different benomyl concentrations. Points represent the mean Optical Density (OD) and error bars represent the standard deviation across three replicates.

### TargetSeeker-MS novel benomyl target discoveries can be confirmed in vitro

GAPDH is among the *H. sapiens* proteins for which a *C. elegans* ortholog was not a predicted as a benomyl target. To our knowledge, GAPDH activity was never previously reported to be affected by benomyl nor was the protein shown to be bound by the compound. We therefore validated the TargetSeeker-MS’ prediction from the Dataset 3 using an orthogonal in vitro approach (see Methods). Figure 5D shows that the enzymatic activity of GAPDH is reduced with an increasing concentration of benomyl, with activity reduced approximately by 50% at 25 uM. This effect confirms the TargetSeeker-MS prediction that GAPDH is bound by benomyl and that this binding affects the in vitro functionality of GAPDH.

## DISCUSSION

### TargetSeeker-MS is compatible with most MS quantification techniques

In this study, spectral counting was used to build protein fractionation profiles to illustrate that TargetSeeker-MS can confidently identify drug targets with simple and cost-effective MS quantitative approaches (label-free quantification). Nevertheless, most types of intensity-based protein quantification strategies (extracted ion chromatogram^24,25^, SILAC^26^, TMT^27^, iTRAQ^28^, etc.) can also be processed by TargetSeeker-MS. In fact, TargetSeeker-MS performance would likely benefit from quantification techniques with improved accuracy. Indeed, spectral counting may not possess the quantification accuracy to enable TargetSeeker-MS to detect slight changes in protein fractionation profiles that are consistent across all replicates. This may particularly be the case for low abundance proteins for which slight changes may be easier to detect with an increase quantification resolution provided by intensity-based quantification techniques.

### TargetSeeker-MS’ applications for therapeutic discovery

EBPS-MS approaches coupled to TargetSeeker-MS can play an important role in the drug discovery process. For instance, high-throughput screenings can allow the identification of a small set of compounds modulating a certain target pathway. These compounds can then be tested against a complex proteome such as the one of a human cell to identify drug bindings and any potential deleterious protein interactions. In the future, EBPS-MS approaches could be applied using TargetSeeker-MS to identify the targets of drugs in induced pluripotent stem cells (iPSCs) and potentially allow the design of better therapeutics.

### Scaling TargetSeeker-MS analysis to multiple compounds

While this study focused on analyzing a single compound in different samples using different EBPS-MS approaches, TargetSeeker-MS’ analysis can be scaled for the analysis of multiple compounds. A given set of untreated samples (four control biological replicates) under the same experimental conditions can be used to identify the drug targets of multiple compounds. Using four controls, TargetSeeker-MS can confidently build its noise model and assess the targets of the tested drugs with two treated biological replicates for a total of six samples, which is the standard number of samples processed in MS-based proteomics experiments comparing two experimental conditions (three replicates from one condition and three replicates from a second condition). Additionally, the noise model built by TargetSeeker-MS can be reused to evaluate supplementary compounds without the analysis of more untreated samples. Since the algorithm only requires the analysis of duplicate drug treated samples, the total number of samples needed to identify the targets of two compounds under the same experimental conditions would be eight, and three compounds would require ten samples (i.e. required samples = 4 + number of compounds × 2). TargetSeeker-MS therefore eases the scalability of the analysis of multiple compounds in contrast to an approach that requires a paired control with each treated sample.

### Drug repositioning applications

The rapidly growing field of drug repositioning depends on the use of techniques like EBPS-MS for the systematic identification of the targets of a drug. The TargetSeeker-MS software package will facilitate the implementation of these techniques and will have a significant impact on the field. Over time the drug discovery pipeline has become increasingly long and costly. Indeed, it takes on average 13.5 years for a drug to reach the market from the start of the investigation^29^. The drug discovery process is also highly failure-prone with a success rate of less than 10%^29^. Drug repositioning is an effort to test whether compounds for which the safety is known and that were approved by organizations such as the FDA could have applications for treatment of disease processes other than the one they were originally designed for. This accelerates the drug discovery process and minimizes the expense and failure risks. Success stories such as duloxetine, which was originally developed to treat depression and is now also used to improve the condition of stress urinary incontinence victims^30^, and crizotinib, which was created to treat anaplastic large-cell lymphoma and was repurposed for the treatment of non-small-cell lung cancer^31^, will hopefully become more common with the increased popularity of EBPS-MS and the use of TargetSeeker-MS.

While EBPS-MS approaches are emerging as fast efficient methods to identify drug targets in complex proteome, no user-friendly approaches have been proposed to assess the confidence of drug binding predictions. We have shown that TargetSeeker-MS can identify high-confidence drug targets with a limited number of control samples. We also demonstrate that TargetSeeker-MS recapitulates predictions of benomyl targets using different EBPS-MS techniques and processing samples from different organisms. In conclusion, our algorithm will favor the growth of the applications of EBPS-MS techniques and improve their impact the field of drug discovery.

## METHODS

### Method overview

We propose a computational approach, TargetSeeker-MS, for the identification of drug targets in energetics-based protein separation (EBPS) coupled to mass spectrometry (MS) datasets. The Bayesian inference approach assesses the confidence that a protein is bound by a given compound. We first describe the two EBPS procedures used in this manuscript (DiffPOP and Thermal shift assay) and the mass spectrometry analysis. We then present TargetSeeker-MS machine learning algorithm. Figure 1 graphically depicts our approach.

### Sample preparation

Mixed stage *C. elegans* were grown and homogenized using a chilled Precellys24 homogenizer (Bertin Instruments) in 80% Phosphoprotein Kit Buffer A (ClonTech, catalog number 635626). Human embryonic kidney cell lysate (HEK293) was prepared from HEK293 cells grown in Dulbecco’s modified Eagle’s medium (D-MEM) with 10% fetal bovine serum (FBS) supplemented with penicillin and streptomycin. Cells were grown (37°C/5% CO_2_) to approximately 80% confluence in tissue culture flasks. Cells were washed twice with DPBS, scrapped from flasks, supplemented with protease inhibitor cocktail (Roche) and lysed by sonication. Protein concentration was determined by BCA assay, lysate was kept at −80°C until use.

*C. elegans* and HEK293 lysate samples were aliquoted and volume adjusted so that they contained five hundred micrograms of proteins in a final volume of 250 μL containing 40 % Phosphoprotein Buffer A (ClonTech). Nineteen aliquots of *C. elegans* lysate were utilized, seven of them were treated with benomyl, while twelve were left untreated (controls). In addition, six aliquots of HEK293 human embryonic kidney cell lysate were utilized, four were left untreated (controls), while two were treated with benomyl (see below).

### Benomyl treatment and DiffPOP protein separation

Lysates of HEK293 and *C. elegans* were incubated with benomyl prior to fractionation. Benomyl (5ul of 10mM stock dissolved in DMSO) was added for the drug treated samples, 5ul of DMSO was added to the control samples. The DiffPOP method was carried out with sequential additions of denaturing solution of 90 % methanol/1% acetic acid (3.75, 8.25, 12.5, 16.25, 20, 42.5, 65, 212.5 and 2000 μL). Each addition was followed by vigorous vortexing and centrifugation (18000 × g for 10 min at 4°C). The supernatant was transferred to a new Eppendorf tube, denaturing solution was added, sample was vortexed and centrifuged. The process was repeated to produce ten pelleted fractions. All resulting pellets were washed with 400 μL ice-cold acetone and centrifuged (18000 × g for 10 min at 4°C). Pellets were air-dried and digested with trypsin (see below).

### Thermal shift assay protein separation

Four *C. elegans* untreated samples and two benomyl treated samples were each separated into 10 fractions using a thermal shift assay procedure slightly modified from the version presented by Savitski et al.^10^ Lysates of *C. elegans* were prepared and incubated with benomyl, as described above, prior to the thermal shift assay. Lysates were heated in an Eppendorf heated shaker block at 20°C for 10 min. Samples were centrifuged at 4°C for 5min. The supernatant was transferred to a new Eppendorf tube and heated to 25°C for 5min. The process was repeated for 30, 35, 40, 50, 60, 75, 90°C to produce 9 pelleted fractions and one supernatant sample. The protein in the final soluble sample was pelleted by addition of methanol/chloroform. All resulting pellets were washed with 400 μL ice-cold acetone and centrifuged (18000 × g for 10 min at 4°C). Pellets were air-dried and digested with trypsin (see below).

### Liquid chromatography coupled to MS/MS analysis

Dried pellets from the DiffPOP and thermal shift assay were dissolved in 8 M urea/100 mM TEAB, pH 8.5. Proteins were reduced with 5 mM tris(2-carboxyethyl)phosphine hydrochloride (TCEP, Sigma-Aldrich) and alkylated with 10 mM chloroacetamide (Sigma-Aldrich). Proteins were digested overnight at 37 °C in 2 M urea/100 mM TEAB, pH 8.5, with trypsin (Promega) at a ratio of 1:100 (enzyme:protein). Digestion was stopped with formic acid (5 % final concentration). Debris was removed by centrifugation. Final volume of each digest was 100ul, and 10ul of each digested fraction was used for analysis by liquid chromatography coupled to tandem mass spectrometry (LC-MS/MS).

The digested samples were analyzed on a Q Exactive mass spectrometer (Thermo Fisher Scientific). The digests were injected directly onto a 2 cm desalting column attached to a 20cm, 100um ID analytical column with pulled tip. Both were packed with 5 μm ODS-AQ C18 resin, (YMC). Samples were separated at a flow rate of 400nl/min on an Easy nLCII (Thermo Fisher Scientific). Buffer A was 5 % acetonitrile and 0.1 % formic acid, buffer B was 80 % acetonitrile and 0.1 % formic acid. The following gradient was utilized: 1-10% B over 5 min, an increase to 45% B over 90 min, an increase to 80% B over another 15 min and held at 80% B for 5 min of washing before returning to 1% B during the final 5 min for a 120 min total run time. Column was re-equilibrated with 10ul of buffer A prior to the injection of sample. Peptides were eluted directly from the tip of the column and nanosprayed directly into the mass spectrometer by application of 2.5kV voltage at the back of the column. The Q Exactive was operated in a data dependent mode. Full MS^1^ scans were collected in the Orbitrap at 70K resolution with a mass range of 400 to 1800 m/z and an AGC target of 1e^6^. A top 10 acquisition method was utilized with HCD fragmentation at 25NCE, resolution of 17.5k, AGC target of 1e^5^ and an underfill ratio of 0.1%. Maximum fill times were set to 60ms and 120ms for MS and MS/MS scans respectively. Quadrupole isolation at 2 m/z was used, singly charged and unassigned charge states were excluded, and dynamic exclusion was used with exclusion duration of 15 sec.

### Peptide and protein identification and quantification

Protein and peptide identification were done with the Integrated Proteomics Pipeline – IP2 (Integrated Proteomics Applications). Tandem mass spectra were extracted from raw files using RawConverter^32^ and searched with ProLuCID^33^ against human SwissProt UniProt^34^ (downloaded on March 25, 2014) and Wormbase^35^ (version WS234) protein sequence databases. The search space included all fully-tryptic and half-tryptic peptide candidates. Carbamidomethylation on cysteine was considered as a static modification. Data was searched with 50 ppm precursor ion tolerance and 600 ppm fragment ion tolerance. Data was filtered to 10 ppm precursor ion tolerance post database search. Identified proteins were filtered using DTASelect^35^ and utilizing a target-decoy database search strategy to control the false discovery rate to 1% at the protein level. To maximize identification specificity only proteins that were identified by at least two different peptides in a given fraction were considered. A single protein identification was retained for the analysis when multiple proteins were inferred from the same peptides across all fractions. Redundant protein identifications were discarded. Proteins were quantified in each fraction by the sum of the spectral counts of their corresponding peptides identified in the fraction.

### Validation of enzymatic activity alteration

A GAPDH activity assay kit (BioVision Inc. Milpitas, CA, Catalog # K680-100) was used to measure GAPDH activity in the presence of benomyl. Benomyl was dissolved in dimethyl sulfoxide (DMSO) and diluted in DMSO to concentrations ranging from 10 mM to 250 uM. The enzyme mixture consisted of 0.5 ul of GAPDH positive control (prepared as instructed by the kit) and 44.5 ul Assay Buffer. The reaction mixture was 2 ul GAPDH Substrate, 2 ul GAPDH Developer and 46 ul Assay Buffer. For each test concentration, 5 ul of drug in DMSO was added followed by 45 ul of enzyme mixture. After a 5 minute incubation at RT, 50 ul of reaction mixture was added. The optical density (OD) at 450 nm was measured at 30 minutes.

### Protein fractionation profiles

Protein fractionation profiles are defined as follows. For a drug treated sample *t*, let a protein *p* be associated with a fractionation profile 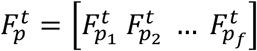, where *f* is the number of fractions the sample is divided into and 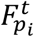, where *i* = 1, …, *f*, is the spectral count of *p* in fraction *i*. 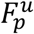 is defined similarly for untreated samples *u*. Fractionation profiles 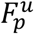 and 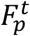 are computed for all *p* ∈ *P*, where *P* is the set of all proteins that were identified with more than 5 spectral counts across all fractions of a sample. Finally, let the fractionation profiles be normalized such that final protein fractionation profiles are defined as follows: 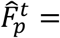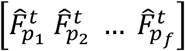, where 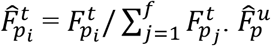 is obtained similarly.

We hypothesize that upon the binding of a drug to a protein, the stability of this protein will be changed. This will affect the fractionation profile of that protein. To measure the change in the fractionation profiles of a protein in the treated and untreated samples we compute the distance between the fractionation profiles of a protein *p* between the drug treated sample 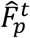 and the untreated samples 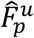 as the Euclidean distance 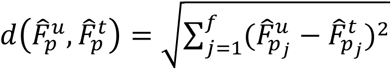. We then compute 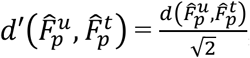, to ensure that the measure falls between 0 and 1 inclusively. We then define the similarity 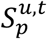 for *p* between two samples’ fractionation profiles 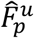 and 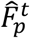 as 1 − 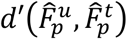.

### Significance assessment of fractionation profile changes

Biological and technical variation may cause fractionation profiles to differ. Such a variation can be captured by computing the similarity 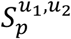 of two fractionation profiles 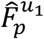 and 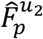 of a protein obtained from two untreated samples *u*_1_ and *u*_2_ under the same experimental conditions. A model specifying for each protein *p* the null distribution of the similarity between untreated protein fractionation profiles could therefore be built. Nevertheless, a large number of untreated samples would be required to estimate this distribution accurately. Processing such large numbers of samples with an EBPS-MS would be lengthy and consume vast amounts of resources. However, we showed that using a small number of untreated samples (e.g. four) and assuming that the proteins with similar similarity average values between the different samples will share a similar null distribution, we can build a noise model of the fractionation profiles by pooling proteins within a given similarity range. Such a model can then allow us to evaluate the significance of the difference of the similarity between a treated sample and the untreated ones. We used an approach inspired by a previously described method used to assess the confidence of protein-protein interactions^36^ to build this model and to assess the significance of the similarity difference between treated and untreated samples. Our novel computational approach is described below.

#### Step 1: Building the similarity noise model from untreated samples

For all untreated sampled that were processed using a EBPS-MS approach TargetSeeker-MS first computes the protein fractionation profiles 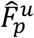 of all identified protein *p*. TargetSeeker-MS then estimates 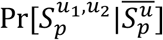, which represents the probability of observing a similarity 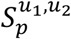 given the true mean similarity of fractionation profiles of *p* in untreated samples 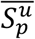. Our algorithm estimates 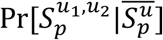 using a leave-one-out scheme on pairs of untreated samples. More specifically, for all untreated sample pairs (*u*_1_, *u*_2_) ∈ *U*×*U*, where *U* is the set of all untreated samples, we compare 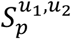 to

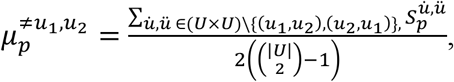

the average of the similarity between all fractionation profiles except those of the sample pair (*u*_1_, *u*_2_). We therefore assume that 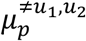 represents a fair approximation of 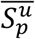. Now let *M* be a 100×100 matrix and *M*(*a*, *b*) represent the frequency where 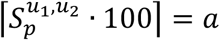 and 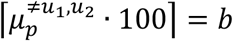. Our goal is to use the frequency matrix *M* to estimate 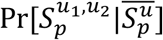, however due to its dimensions, an important number of entries in *M* have a zero value. This results in an estimator for 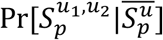, which would often yield probabilities that are equal to zero. In order to solve this problem, we used a *k*-nearest neighbor smoothing algorithm we have previously described^36^. to build the smoothed frequency matrix *M*′. Briefly, let *N*_*δ*_ (*a*, *b*) = {*a*′, *b*′):|*a* − *a*′| ≤ *δ*, |*b* − *b*′| ≤ *δ*} be the set of cells surrounding *M*(*a*, *b*) with a distance *δ*. For each entry in *M*, *δ* is computed such that 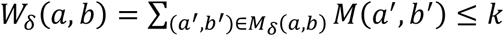 and 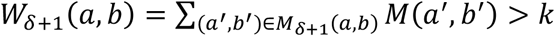. The smoothed matrix *M*′ is therefore computed as following:

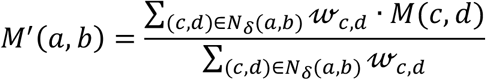

where

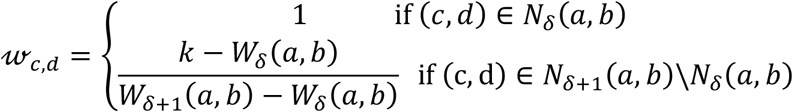

Upon the empirical analysis from a large range of *k* values, *k* = 20 was determined to yield a good balance of smoothing, while conserving the original signal of the data and therefore providing the best results. Therefore, 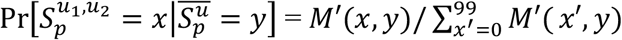 provides us a good estimate of the noise of the similarity between two fractionation profiles.

#### Step 2: Posterior distribution of the mean similarity between fractionation profiles from untreated samples

Using Bayes’ rule and assuming the conditional independence of the similarity observations given their true means, the posterior distribution of 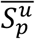 is computed as follows:

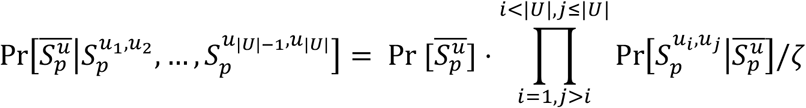

where *ζ* is a normalizing constant, and 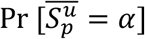 corresponds to the fraction of proteins with 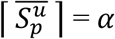. This calculation provides us the distribution of the mean similarity of the fractionation profile of a protein between untreated samples given the set of similarity observations

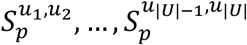

#### Step 3: Significance assessment

The goal of TargetSeeker-MS is to assess the significance of the change in the fractionation profile of each protein in untreated samples and a drug treated sample. In order to do so, we first compute the average of the similarity values between 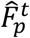 and 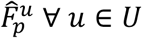 as follows, 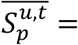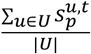. We then assess the significance of the change in similarity for the drug treated fractionation profile for each protein by computing a *p*-value using our noise model of the similarity of the fractionation profiles of a protein in untreated samples (computed in *Step 2*):

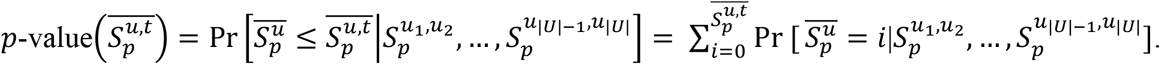

#### Step 4: False discovery rate estimation

The *p*-values computed in *Step 3* may be underestimated as a result of the possible violation of some of the assumptions made when building our noise model. To assess the specificity of its predictions, TargetSeeker-MS uses a hypothesis-free approach to compute a False Discovery Rate (FDR) for each protein. Given a *p*-value threshold *p*, TargetSeeker-MS computes *FDR*(*p*), which represents the fraction of false positive predicted drug binders that obtained a *p*-value < *p*. *FDR*(p) is computed using a leave-one-out strategy, where each untreated biological replicate is alternately left out from the rest of the untreated samples for the *p*-value calculation and considered as if it was a treated sample. Specifically, ∀ *u*′ ∈ *U*, TargetSeeker-MS computes *p*-value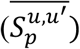 for all proteins *p* quantified in the untreated sample *u*′ (left out) and *p*-value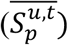 for all proteins *p* quantified in the treated sample *t* using the set of untreated samples excluding *u*′ to build the noise model. TargetSeeker-MS then computes for the FDR for the *p*-value *p* as follows:

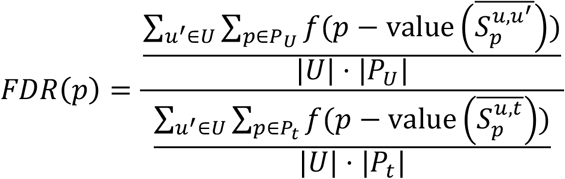

where

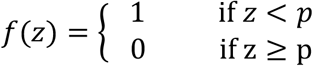

and *P*_*U*_ and *P*_*t*_ are the sets of proteins with fractionation profiles in the untreated and treated samples respectively. TargetSeeker-MS’ algorithm design, involving two independent leave-one-out schemes, therefore requires a minimum of four untreated sample replicates to allow the confidence assessment of putative drug targets.

#### Step 5: Biological replicates of drug treated samples

To maximize the stringency of its drug target predictions when given biological replicates of drug treated samples, TargetSeeker-MS allows the user to analyze both replicates independently and to report the proteins that are considered high-confidence drug targets in all replicates. Alternatively, TargetSeeker-MS can also analyze all biological replicates of drug treated samples simultaneously. In this context, TargetSeeker-MS computes the average of the similarity values between 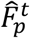 and 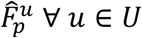, *t* ∈ *T*, such that 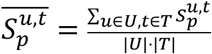, where *T* is the set of treated samples. The two approaches tend to yield similar results with the former being slightly more stringent (see Results).

### Computation of fold-change of similarity difference

To maximize the specificity of TargetSeeker-MS’ predictions, putative drug targets must be associated with a FDR < 0.10, but a Fold-change of Similarity Difference (FSD) above a given threshold can also be used. The FSD of a protein *p* is calculated as follows:

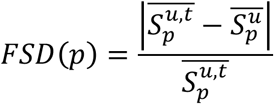

### Implementation and availability

TargetSeeker-MS is implemented as a web-based fast Java program that is available at this address: http://targetseeker.scripps.edu/. Example input and output files are provided online. Given a set of untreated and drug treated samples that were fractionated using an EBPS approach, TargetSeeker-MS computes a FDR for all proteins with a fractionation profile in both conditions. Note that intensity-based protein quantification can also be provided as input to TargetSeeker-MS. The methods are also implemented as stand-alone, open-source, platform independent, command-line-based Java program, which is available at this address http://targetseeker.scripps.edu/files/ and on GitHub: https://github.com/proteomicsyates. The mass spectrometry proteomics data have been deposited to the ProteomeXchange Consortium via the PRIDE partner repository with the dataset identifier PXD010799. Data are stored in Pride Archive. To access the data files, please visit https://www.ebi.ac.uk/pride/archive/login and use the following user name: and password: 4nyA0clB.

### Alternative approaches

#### Z-score filtering

We developed an alternative method to benchmark the TargetSeeker-MS algorithm. The *Z*-score method computes a Z-score for each protein *p* by comparing the similarity of its fractionation profiles between the treated and untreated samples 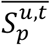 to the mean 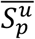 and the standard deviation 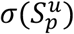 of the similarity of the fractionation profiles in untreated samples: 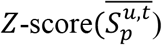 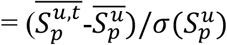. This alternative approach may outperform the TargetSeeker-MS algorithm if the variance of the fractionation profile similarity values in untreated samples of proteins with close mean similarity values differs significantly from protein to protein. This is due to the pooling of similarity values from different proteins in TargetSeeker-MS, which makes the assumption that this variation is low. Nevertheless, since no data pooling is performed with the *Z*-score method, the estimation of the variance is performed using a small number of values (i.e. the total number of untreated sample pairs). Furthermore, the *Z*-score method assumes that the noise of the fractionation profile similarity values in untreated samples is normally distributed, which cannot be unequivocally verified. FDR values for all *Z*-scores were estimated using the same leave-one out approach as described in *Step 4*.

#### Savitski et al. statistical approach

We implemented the statistical approach adapted from the article from Cox et al.^11^ We analyzed all EBPS-MS datasets (Dataset 1, 2 and 3) with this implementation in the fashion it was applied in the Savitski et al.’s article^10^, with the only difference that melting curve slopes were replaced with similarity values. It should be noted that this modification does not affect the validity of the statistical approach nor does it change any of its assumptions about input values. Due to the different nature of the algorithm, FDR values were estimated with a slightly modified procedure than the one described in *Step 4*. The algorithm was fed the average similarity values of all proteins *p* ∈ *P*_*u*_ for which a fractionation profile was computed in untreated samples to which *p*-values were associated. *p*-values were then calculated for all proteins *p* ∈ *P*_/_ for which a fractionation profile was computed based on the average similarity values of these profiles between treated and untreated samples. FDRs were then associated to each *p*-value *p* as follows:

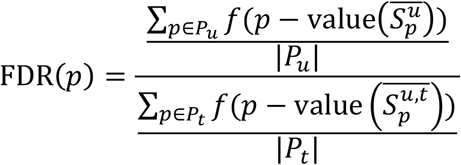

where

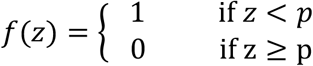

### Gene Ontology enrichment analysis

To investigate the mechanism of action of benomyl, we evaluated the statistical enrichment of Gene Ontology terms^37^ among the proteins predicted as its targets by TargetSeeker-MS in the three different datasets using Ontologizer^38^. We tested the enrichment of molecular functions, biological processes, and cellular components (with the complete set of proteins associated with a fractionation profile as background). Ontologizer uses a modified Fisher’s exact test to assess the statistical significance of the enrichment of Gene Ontology terms and the Bonferroni correction to correct for multiple hypothesis testing^38^.

### Protein ortholog determination

Orthologous protein targets between *H. sapiens* and *C. elegans* were determined using the Blastp algorithm^39^. Proteins associated with a sequence identity between the two species > 50% and an E-value < 10^−10^ were considered orthologs.

## ACKNOWLEDGEMENTS

The authors are grateful to Claire M. Delahunty for helpful discussions and comments. They acknowledge funding from the following National Institute of Health grants: P41 GM103533, R01 MH067880, R01 MH100175, UCLA/NHLBI Proteomics Centers (HHSN268201000035C), and U54GM114833 to J.R.Y and NSERC Discovery grant to M.L.A. M.L.A. held a postdoctoral fellowship from the Fonds de recherche du Québec – nature et technologies (FRQNT).

## AUTHOR CONTRIBUTIONS

M.L.A. designed the computational approach, which was implemented by A.P. with help from M.L.A. and S.M.B. The DiffPOP method was developed by J.K.D., A.F.M.P. and J.J.M. The TSA approach was adapted by J.K.D. The DiffPOP, TSA, and LC-MS/MS procedures were performed by J.K.D., A.P. and W.L. The enzymatic assay was performed by J.J.M. C. elegans sample processing was performed by M.P. Finally, M.L.A., A.P., J.K.D., J.J.M. and J.R.Y. wrote the manuscript.

## SUPPLEMENTARY FIGURES

**Supplementary Figure S1.** Graphical depiction of the datasets presented in this study.

**Supplementary Figure S2.** Heatmap representation of the similarity between protein fractionation profiles for all pairs of untreated samples of the TSA/*C. elegans* dataset. All proteins with a sufficient abundance to compute a fractionation profile in all untreated samples are displayed.

**Supplementary Figure S3.** Heatmap representation of the similarity between protein fractionation profiles for all pairs of untreated samples of the DiffPOP/HEK293 dataset. All proteins with a sufficient abundance to compute a fractionation profile in all untreated samples are displayed.

**Supplementary Figure S4.** Three-dimensional plot representation of the probability matrix of 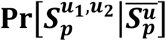 based on the smoothed frequency matrix *M*′ for four untreated samples from the DiffPOP/*C. elegans* dataset.

**Supplementary Figure S5.** Heatmap representation of the similarity between protein fractionation profiles for all pairs of benomyl treated samples of the DiffPOP/*C. elegans* dataset. All proteins with a sufficient abundance to compute a fractionation profile in all benomyl treated samples are displayed.

**Supplementary Figure S6.** Heatmap representation of the similarity between protein fractionation profiles for the two benomyl treated samples of the TSA/*C. elegans* dataset. All proteins with a sufficient abundance to compute a fractionation profile in both benomyl treated samples are displayed.

**Supplementary Figure S7.** Heatmap representation of the similarity between protein fractionation profiles for the two benomyl treated samples of the DiffPOP/HEK293 dataset. All proteins with a sufficient abundance to compute a fractionation profile in both benomyl treated samples are displayed.

**Supplementary Figure S8.** Distributions of the fractionation profile similarity values in both untreated and benomyl treated samples of the TSA/*C. elegans* dataset.

**Supplementary Figure S9.** Distributions of the fractionation profile similarity values in both untreated and benomyl treated samples of the DiffPOP/HEK293 dataset.

## SUPPLEMENTARY TABLES

**Supplementary Table S1:** *C. elegans* proteins identified through DiffPOP-MS analysis along with their associated statistics as calculated by TargetSeeker-MS when combining all treated samples.

**Supplementary Table S2:** Benomyl treated sample 1 *C. elegans* proteins identified through DiffPOP-MS analysis along with their associated statistics as calculated by TargetSeeker-MS.

**Supplementary Table S3:** Benomyl treated sample 2 *C. elegans* proteins identified through DiffPOP-MS analysis along with their associated statistics as calculated by TargetSeeker-MS.

**Supplementary Table S4:** Benomyl treated sample 3 *C. elegans* proteins identified through DiffPOP-MS analysis along with their associated statistics as calculated by TargetSeeker-MS.

**Supplementary Table S5:** *C. elegans* proteins identified as benomyl targets in each treated sample processed with DiffPOP-MS.

**Supplementary Table S6:** Gene Ontology enrichment analysis complete results for the benomyl targets identified in the DiffPOP-MS-*C. elegans* dataset.

**Supplementary Table S7:** *C. elegans* proteins identified through Thermal Shift Assay-MS analysis along with their associated statistics as calculated by TargetSeeker-MS when combining all treated samples.

**Supplementary Table S8:** Benomyl-treated sample 1 *C. elegans* proteins identified through Thermal Shift Assay-MS analysis along with their associated statistics as calculated by TargetSeeker-MS.

**Supplementary Table S9:** Benomyl-treated sample 2 *C. elegans* proteins identified through Thermal Shift Assay-MS analysis along with their associated statistics as calculated by TargetSeeker-MS.

**Supplementary Table S10:** *C. elegans* proteins identified as benomyl targets in both treated samples processed with TSA-MS.

**Supplementary Table S11:** Gene Ontology enrichment analysis complete results for the benomyl targets identified in the TSA/*C. elegans* dataset.

**Supplementary Table S12:** HEK 293 proteins identified through DiffPOP-MS analysis along with their associated statistics as calculated by TargetSeeker-MS when combining both treated samples.

**Supplementary Table S13:** Benomyl-treated sample 1 HEK 293 proteins identified through DiffPOP-MS analysis along with their associated statistics as calculated by TargetSeeker-MS.

**Supplementary Table S14:** Benomyl-treated sample 2 HEK 293 proteins identified through DiffPOP-MS analysis along with their associated statistics as calculated by TargetSeeker-MS.

**Supplementary Table S15:** HEK 293 proteins identified as benomyl targets in both treated samples processed with DiffPOP-MS.

**Supplementary Table S16:** Gene Ontology enrichment analysis complete results for the benomyl targets identified in the DiffPOP/HEK 293 dataset.

## SOURCE DATA

**Figure 2A** - **Source Data:** Csv file containing the data presented in the heatmap of Figure 2A.

**Figure 2B** - **Source Data:** Excel file containing the data of the distributions presented in Figure 2B.

**Figure 2C-D** - **Source Data:** Excel file containing the data of the distributions and protein fractionation profiles presented in Figures 2C and 2D.

**Figure 3A** - **Source Data:** Excel file containing the data of the benchmarking presented in Figure 3A.

**Figure 4A** - **Source Data:** Excel file containing the data of the benchmarking presented in Figure 4A.

**Figure 5A** - **Source Data:** Excel file containing the data of the benchmarking presented in Figure 5A.

**Figure 5D** - **Source Data:** Excel file containing the data of the results of the GAPDH activity assay presented in Figure 5D.

